# Genetic diversity and differentiation of a mycoheterotrophic orchid (*Cymbidium macrorhizon)* under urbanization

**DOI:** 10.1101/2024.09.01.610666

**Authors:** Naohiro I. Ishii, Satoshi Yamamoto, Yuki Iwachido, Kei Uchida

## Abstract

Urbanization exerts substantial pressures on genetic diversity of plant species. However, depending on species-specific life history, the direction/magnitude of urbanization impacts can vary. To elucidate relationships between life history and urbanization effects, there are needs to accumulate the knowledge on genetic diversity/differentiation along urban-rural gradients for species with unique traits. We examined these facets based on genome-wide single nucleotide polymorphisms of a mycoheterotrophic and vegetative-dormant orchid, *Cymbidium macrorhizon*, for eleven populations in remnant forests along an urban gradient within the Tokyo metropolitan area, Japan. The reduced inbreeding coefficient and increased genetic differentiation were observed with increased proportion of surrounding urban land-use 50 years ago rather than in recent years. This pattern might reflect lowest heterozygosity under intensive urbanization due to population bottleneck and genetic drift due to habitat shrinkage and fragmentation. The significant impacts of past landscape on the indices might indicate time lags of genetic erosion, namely intra-specific extinction debt, due to longevity and dormancy. Therefore, 30% increase of urban land-use since 1970s has not yet affected genetic erosion, resulting in its progression in the future. We emphasize the importance not only to assess genetic diversity but also to connect the assessments with life history and spatiotemporal urbanization impacts.

## Introduction

High human pressures have caused shrinkage, fragmentation, and changing quality of habitats, thereby leading to drastic changes in species and genetic diversity (Grimm et al. 2008; Seto et al. 2012; Fukano et al. 2023). One of the significant human pressures on habitats is urbanization, which can alter habitat quality and/or quantity of habitats and may cause reduced genetic diversity, and the increase of inbreeding and genetic differentiation among populations by affecting population size and gene flow (Johnson and Munshi-South 2017; Miles et al. 2019). The patterns of gene flow (limitation/facilitation) can vary depending on the focal species under evaluation for genetic diversity, reflecting their functional traits such as long-term life history traits (Miles et al. 2019).

Urban ecosystems are even hotspots for plant diversity as well as animal diversity compared to other ecosystems (Ives et al. 2016; Kasada et al. 2017; Theodorou et al. 2017; Uchida et al. 2023). Indeed, plant species in urban areas largely contribute to ecosystem functions and services (Schwarz et al. 2017; Theodorou et al. 2020). The effective conservation of plant diversity and its ecosystem services requires the accumulation of information on genetic diversity, which is associated with the reduction in adaptive potential to environmental changes and reproductive fitness in members of urban plant communities (Reed and Frankham 2003; O’Grady et al. 2006; Willi et al. 2006; Losdat et al. 2014).

However, most previous studies focused on the differences in dispersal ability related to functional traits that contributed to urbanization impacts on animals (mainly mammals, arthropods, and birds) (Miles et al. 2018; Schmidt et al. 2020), little is known about the genetic erosion and contemporary evolution of plant species due to urbanization in remnant habitats compared to in highly urbanized land uses (Dubois and Cheptou 2017; Yakub and Tiffin 2017; Johnson et al. 2018). Although genetic erosion in urban areas generally occurs in island-like habitats with reductions in area and fragmentations (Honnay and Jacquemyn 2007), the intensity of genetic erosion is likely to be variable depending on life history traits such as longevity, formation of seed banks, and seed/vegetative dormancy (Fuller and Doyle 2018; Figueiredo et al. 2019).

In addition, individual-, population-, and meta-population-level dynamics relating to species-specific life history traits can also lead to the occurrences of time lags in genetic erosion and evolution after the reduction in habitat area (Helm et al. 2009; Plue et al. 2017; Reisch et al. 2017; Fuller and Doyle 2018; Aavik et al. 2019). These time lags in urban remnant ecosystems have not been quantified, which points to the possibility of underestimations of urbanization impacts on genetic erosion or contemporary evolvability. In particular, the quantification of genetic erosion and its time lag can implicate that the payment of the population-level extinction debts can accelerate in the future (Figueiredo et al. 2019). Therefore, we should provide important information for conservation prioritization and strategies to maintain and enhance genetic diversity.

Orchidaceae is one of the most vulnerable family to urbanization, and their extinctions in urban areas have been reported worldwide (Duncan et al. 2011; Vogt-Schilb et al. 2015; Khapugin et al. 2020). This family includes species with a wide range of life history traits such as mycoheterotrophy, long lifespan, and seed/vegetative dormancy (Shefferson 2009; Shefferson et al. 2020). The interactions between orchids and fungi exhibit a higher level of mycotrophic specialization than that in other taxa (Brundrett 2004). Considering mycorrhizal specialization to host tree species, the sparse remnant forests in urban cities can lead to the high rarity of orchids, resulting in the promotion of genetic erosion by urbanizations. On the other hand, long lifespan and dormancy can give rise to generation overlaps, which can maintain or increase genetic diversity under selective pressures (Ellner and Hairston 1994; Ellner 1996; Tsuzuki et al. 2022). However, the urbanization impacts on the mycoheterotrophic species with the traits of long lifespan and vegetative dormancy are unclear. The assessment of urbanization impacts and their time lag in these species across urban ecosystems helps decision making in conservation prioritization and urgency of conservation implementation.

Here, the present study addressed the relationships between genetic diversity/differentiation of *Cymbidium macrorhizon* Lindl. and changes in landscape elements (forests and urban land-uses) in remnant forest areas in a metropolitan area of Japan. *C. macrorhizon* has a distinctive life history such as mycoheterotrophy, autonomous self-pollination, and low frequency of pollination (Motomura et al. 2010; Suetsugu 2014). Based on the dataset of *de novo* single nucleotide polymorphisms (SNPs), we evaluated genetic diversity and differentiation of this species in multiple remnant forests across the Tokyo major metropolitan area (MMA). To identify (1) the influences of landscape variables (the proportion of urbanized and forest area) on genetic indices, (2) the relative contribution of current and past (approx. 1970s) patterns of the landscape elements.

## Materials and Methods

### Study species and sites

The present study focused on the genetic diversity of *C. macrorhizon* across remnant forests in the Tokyo MMA, Japan (Figure 1). *C. macrorhizon* is a leafless and rootless mycoheterotrophic orchid that is distributed widely in subtropical to warm-temperate Asia across the Himalayas, and from Indochina to Japan (Yukawa 2015). This species occurs in evergreen or deciduous broadleaf forests, and mixed pine forests in Japan (Ogura-Tsujita et al. 2012; Suetsugu 2014; Suetsugu et al. 2018). In the Tokyo MMA, the habitat of *C. macrorhizon* is the forest floor of deciduous broadleaf forests (dominated by *Quercus serrata* Murray and *Q. acutissima* Carruth.). The nectarless flowers of *C. macrorhizon*, which have the structure for autonomous self-pollination, are pollinated by insects (mainly honeybees) at low frequency (Suetsugu 2014). The dust-like seeds are assumed to be dispersed by wind as with most orchid species (Eriksson and Kainulainen 2011). The chromosome number of *C. macrorhizon* is 2n = 38 (Pal et al. 2020). *C. macrorhizon* is listed in the Japanese Red List as Vulnerable (VU) (Japanese Ministry of the Environment 2020) due to population decline caused by habitat shrinkages associated with deforestations and/or afforestation.

**Figure 1.**
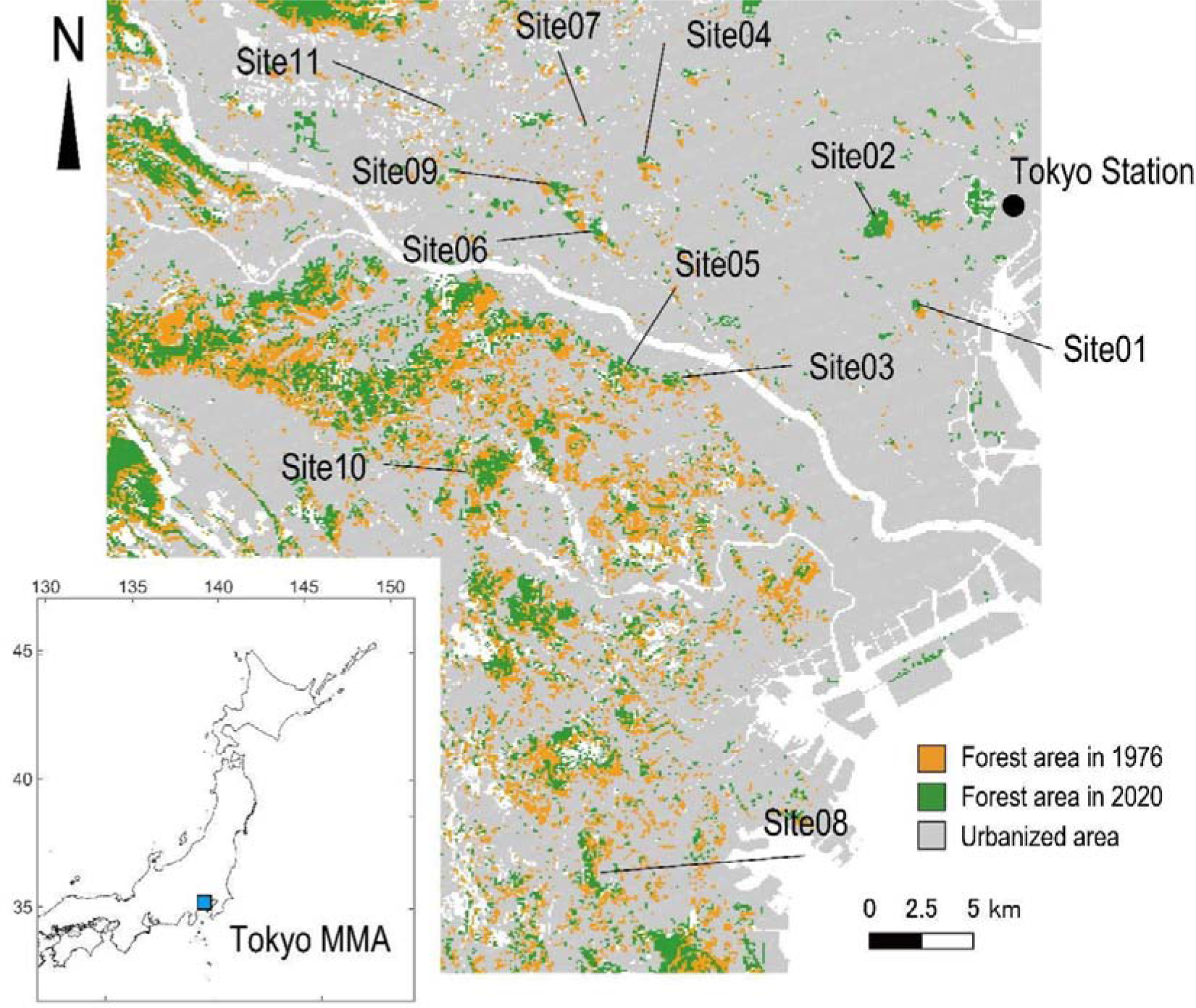
Spatial distribution of study sites across the Tokyo major metropolitan area (MMA). The urbanized area (grey), and forest area of 1976 (orange) and 2020 (green) were denoted.

We selected 11 remnant forests with different proportions of surrounding urbanized area and forest area across the Tokyo MMA (Figure 1, and Table S1). For each study site, the petal samples of 7–16 individuals (total 126 individuals) were collected from June to October 2020 (Table 1). All samples were placed in silica gel and stored at −20 °C before DNA extraction.

**Table 1.** Number of analyzed individuals and genetic statistics of 11 populations of *Cymbidium macrorhizon*. *H*_O_ and *H*_E_, average observed and expected heterozygosity; *F*_IS_, inbreeding coefficient; average pairwise *F*_ST_, average value of pairwise fixation index between all pair of each site.

| Site ID | No. of individuals | No. of analyzed individuals | No. of analyzed samples per locus | No. of polymorphic sites | Proportion of polymorphic sites (%) | $H_O$ | $H_E$ | $F_{IS}$ | Average pairwise $F_{ST}$ |
| --- | --- | --- | --- | --- | --- | --- | --- | --- | --- |
| 01 | 8 | 8 | 7.0 | 189 | 0.0036 | 0.225 | 0.168 | -0.084 | 0.171 |
| 02 | 8 | 8 | 7.1 | 198 | 0.0038 | 0.223 | 0.168 | -0.077 | 0.277 |
| 03 | 16 | 13 | 8.5 | 271 | 0.0052 | 0.211 | 0.243 | 0.112 | 0.079 |
| 04 | 14 | 12 | 8.3 | 267 | 0.0051 | 0.185 | 0.201 | 0.079 | 0.074 |
| 05 | 7 | 6 | 4.1 | 223 | 0.0043 | 0.238 | 0.196 | 0.008 | 0.070 |
| 06 | 16 | 15 | 11.5 | 310 | 0.0060 | 0.243 | 0.248 | 0.081 | 0.102 |
| 07 | 8 | 7 | 4.5 | 182 | 0.0035 | 0.199 | 0.178 | 0.021 | 0.105 |
| 08 | 14 | 12 | 9.1 | 239 | 0.0046 | 0.160 | 0.181 | 0.069 | 0.100 |
| 09 | 16 | 15 | 11.2 | 277 | 0.0053 | 0.247 | 0.226 | 0.030 | 0.149 |
| 10 | 8 | 8 | 6.0 | 216 | 0.0042 | 0.171 | 0.171 | 0.029 | 0.088 |
| 11 | 11 | 11 | 7.7 | 238 | 0.0046 | 0.180 | 0.207 | 0.095 | 0.076 |

### DNA extraction, library preparation, and sequencing

To extract total genomic DNA from the petal samples, we first disassembled each sample in a 1.5-mL tube using autoclaved tweezers. Genomic DNA was extracted using the DNeasy Plant Mini kit (Qiagen, Hilden, Germany).

A double-digest restriction site-associated DNA (ddRAD) library (Peterson et al. 2012) was prepared using the Peterson’s protocol with modifications. First, we digested 100 ng of genomic DNA with PstI and EcoRI (New England Biolabs, Ipswich, MA, USA) at 37°C for an hour in a 15 µl volume, containing 10 units of enzyme for each PstI and EcoRI, 2 µl of Cut Smart Buffer (New England Biolabs) and 2.8 µl of deionized water. Then, adapters were ligated at 23°C for 20 min in an 18 µl volume, containing 1 µl of 100 nM adapters that match to sticky ends generated by PstI and EcoRI, 0.2 µl of 100 mM rATP (Promega, Madison, WA, USA), 10,000 units of T4 DNA ligase (New England Biolabs), 0.3 µl of Cut Smart Buffer, and 0.3 µl of deionized water. The reaction solution was then purified with NucleoMag NGS Clean-up and Size Selection kit (Macherey-Nagel, Germany). Next, 16 µl of purified DNA was used in the PCR amplification containing 8 µl of primer solution that includes each 10 µM of forward and indexed-reverse primers and 24 µl of KAPA HiFi HotStart ReadyMix (KAPA Biosystems, Wilmington, MA, USA). Thermal cycling was initiated with a 98°C step for 3 min, followed by 15 cycles of 98°C for 20 sec, 72°C for 1 min, and 72°C for 5 min. The PCR products were purified again with NucleoMag and pooled. The library was sequenced with 150 bp pair-end reads by an Illumina HiSeqX (Illumina, San Diego, CA, USA).

### Read assembling and SNP calling

One base at each end of the processed reads were trimmed and the reads were filtered based on the quality of reads and the adaptor sequences with the default parameter settings using fastp v0.20.1 (Chen et al. 2018). Therefore, we performed the subsequent analysis using the filtered reads at length of 145 bp. The samples with less than 0.1 million reads were removed for analysis. All sequence data were deposited at the DDBJ Sequence Read Archive (DRA) with accession number DRA.

Stacks v2.52 (Catchen et al. 2011) was used to process the filtered reads for *de novo* assembly. We performed the *de novo* assembly without mapping to genome using the function ‘ustacks’ and ‘cstacks’ in Stacks with the default parameter settings: the minimum number of identical reads required to create a stack, m = 3; the nucleotide mismatches between loci within a single individual, M = 2; and the mismatches between loci when building the catalog, n = 1. SNP calling was conducted using the function ‘population’ in Stacks as in the former with one modification: minimum sharing rate of a single SNP among all analyzed individuals, -R = 0.05 for phylogenetic analysis and 0.6 for calculation of genetic indices. Finally, we removed the samples that showed a missing rate of called SNPs higher than 60 % at the latter setting (-R = 0.6). 115 samples remained and were used for the subsequent analysis (Table 1).

### Genetic diversity and differentiation

Maximum likelihood phylogenies were inferred using RAxML v8.2.12 (Stamatakis 2014). The resolution of phylogenetic analysis tends to be higher when the phylogenetic tree is constructed by a larger number of SNPs even with low genotyping rates (Wagner et al. 2013). Thus, we used the dataset with the setting of -R = 0.05 in the function ‘population’ in Stacks. We selected a GTR+CAT model and performed 1000 replicates of parallelized tree search bootstrapping. To detect population genetic structure of *C. macrorhizon* across the Tokyo MMA, filtered super network based on 1000 replicate trees was estimated with the threshold of 500 trees using SplitsTree v4.17.1 (Huson and Bryant 2006). In subsequent analyses, the data set with the setting of -R = 0.6 in the function ‘population’ in Stacks was used. To find SNPs associated with urbanized gradient (i.e., SNPs adaptive to urbanization), we performed an outlier test that can account for the proportional gradient of urbanized area using BayeScEnv v1.1 with the default parameter settings (de Villemereuil and Gaggiotti 2015).

Observed and expected heterozygosity indices (*H*_O_ and *H*_E_) as the indices of genetic diversity within each study site, and inbreeding coefficient (*F*_IS_) were calculated using the function ‘population’ in Stacks. As the index of genetic differentiation among populations we studied, we calculated pairwise *F*_ST_ values using the function ‘pairwise.neifst’ in the R package hierfstat (Goudet 2005) and averaged the values of pairwise *F*_ST_ based on all, neutral, and adaptive SNPs for each study site. To detect genetic differentiation at the population-level, a neighbor net was estimated using SplitsTree v4.17.1 (Huson and Bryant 2006). In the case where there is a distinct population genetic structure, multiple clusters can be split by reticulations. To visualize the spatial patterns of the genetic indices, the inverse distance weighted spatial interpolations based on the genetic indices were conducted using the function ‘IDW (inverse distance weighted)’ in QGIS. To test if there is a significant spatial autocorrelation of pairwise *F*_ST_, we performed the Monte Carlo test of global spatial structures based on Moran’s eigenvector map (MEM) using the function ‘global.rtest’ in the R package adespatial. This test was done for the principal coordinates based on the matrix of pairwise *F*_ST_ and the Delaunay triangles based on coordinate of each study site as a connection network.

### Data analysis

The present study focused on detecting relationships between genetic indices and landscape elements due to urbanization. As landscape variables associated with urbanization, the proportion of surrounding urban land–use area and forest area within a radius of 1000, 3000, and 5000 m was calculated. These values were based on the 100 m mesh land-use data of 1976 and 2020 provided by the Geospatial Information Authority of Japan (https://nlftp.mlit.go.jp/index.html). Firstly, we calculated Moran’s *I* and its significance of genetic indices and landscape variables based on the Delaunay triangles from the coordinates of the study sites to check spatial autocorrelations using the function ‘moran.test’ in the R package spdep. To distinguish which the proportion of (1) urban or forest area within (2) a radius of 1000, 3000, or 5000 m in (3) 1976 or 2020 is more strongly associated with genetic diversity (*H*_E_), inbreeding coefficient (*F*_IS_), and genetic differentiation (average pairwise *F*_ST_ based on all, neutral, adaptive SNPs), we performed the regression analysis by generalized linear models (GLMs) with model selection procedure based on Akaike information criterion (AIC). The response variables were each genetic index (gaussian distribution), while explanatory variables included a single landscape variable and the average number of samples per a SNP site (Table 1). The latter variable, which was common across all models, was included to correct for bias in genetic indices due to the number of individuals analyzed. In other words, the differences in AIC among models imply those in explanatory power (model fit) of each landscape variable.

## Results

The mean number of raw and filtered genomic reads per sample was 3,734,302 and 3,236,296, respectively. The number of monomorphic and polymorphic sites was 5,210,326 (16,989 loci) and 372 (174 loci), respectively. The number of SNPs within each study site ranged from 182 to 277 (Table 1). The average polymorphic rate (SNP rate to monomorphic and SNP sites) was 0.005 %.

The inferred phylogenetic tree showed a radial network topology, which reflected the lack of a distinct population genetic structure of *C. macrorhizon* across the Tokyo MMA (Figure S1). Despite distant geographic separation among study sites, there was no substantial decrease in genetic relatedness between individuals.

The number of neutral and adaptive SNPs was 307 and 65, respectively. *H*_O_, *H*_E_, and *F*_IS_ within each study site ranged from 0.160 to 0.247, from 0.168 to 0.248, and from −0.084 to 0.112, respectively (Table 1). Only the population of the Site01 and Site02 had the negative values of *F*_IS_. The average of pairwise *F*_ST_ based on all SNPs ranged from 0.070 to 0.277 (Table 1). The average values of pairwise *F*_ST_ based on neutral and adaptive SNPs were ranged from 0.021 to 0.109 and from 0.398 to 0.903, respectively. In the Site01 and Site02, which had high proportion of urban area in the past, those populations showed the lowest *H*_E_ and *F*_IS_, and the highest values of average pairwise *F*_ST_ (Figure 2). The neighbor net using pairwise *F*_ST_ based on all SNPs between the study sites showed that the populations of the Site01, Site02, and Site09 had the different genetic composition from the others (Figure S2), which was different from the individual-level phylogenetic relationship across the study sites (Figure S1). The significant pattern of spatial autocorrelation in genetic differentiation (pairwise *F*_ST_) was not detected as the result of the test based on MEM (Monte Carlo test, *P* = 0.483).

**Figure 2.**
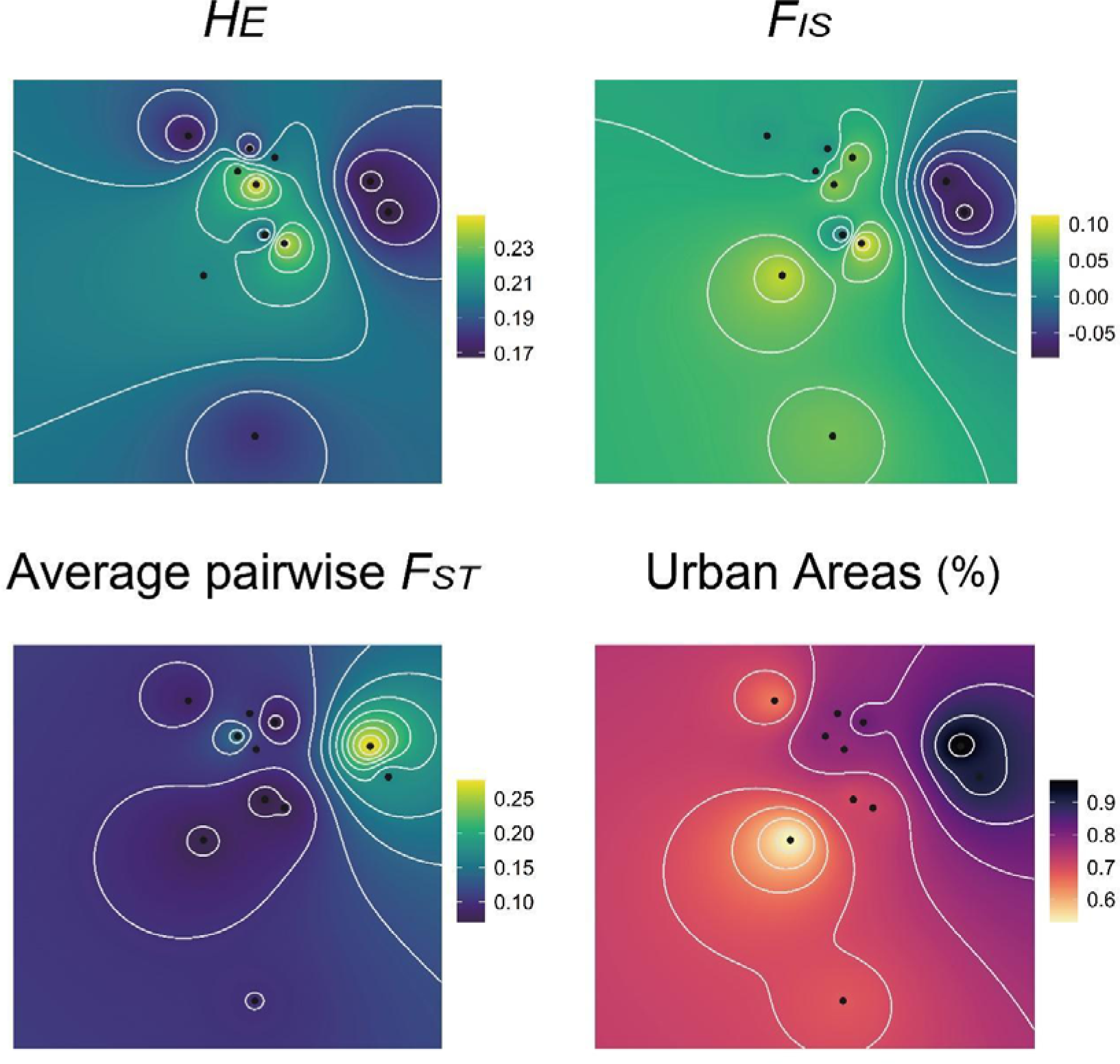
Results of spatial interpolation based on the genetic indices and the proportion of urban area. *H*_E_, average expected heterozygosity; *F*_IS_, inbreeding coefficient; average pairwise *F*_ST_, average value of pairwise fixation index between all pair of each site.

The result from GLM showed that *H*_E_ was not significantly related to the landscape variables (urban land-use and forest area) within any radius in any period (Table 2, and Figure 3). We detected the significantly positive and negative relationships between *F*_IS_, and urban/forest area, respectively (Figure 3). Especially, the model including the proportion of urbanized area in 1976 showed the lower level of AIC than the others (Table 2). The model with the proportion of urban area within 5000m in 1976 showed the highest model fitting, and its regression coefficient β was −0.388 (95 % CI: −0.595 – −0.181). Average pairwise *F*_ST_ based on all SNPs was significantly positively correlated to the proportion of urban area (Figure 3). The best fit models for average pairwise *F*_ST_ based on all SNPs included the proportion of urban area within 3000 and 5000 m in 1976 (β = 0.292 and 0.385, respectively; Table 2). In addition, as the result of GLM for average pairwise *F*_ST_ based on neutral and adaptive SNPs, the significant positive correlations between both indices and the proportion of urbanized area within 5000 m in 1976 (β = 0.183 and 0.772, respectively; Table S2, and Figure S3). Note that all the genetic indices did not show significant spatial autocorrelations (Table S2). On the other hand, the significant and positive spatial autocorrelations in some of the landscape variables (urbanized rates within 2000 and 3000 m, and forest rate within 3000 m) were observed (Table S2).

**Figure 3.**
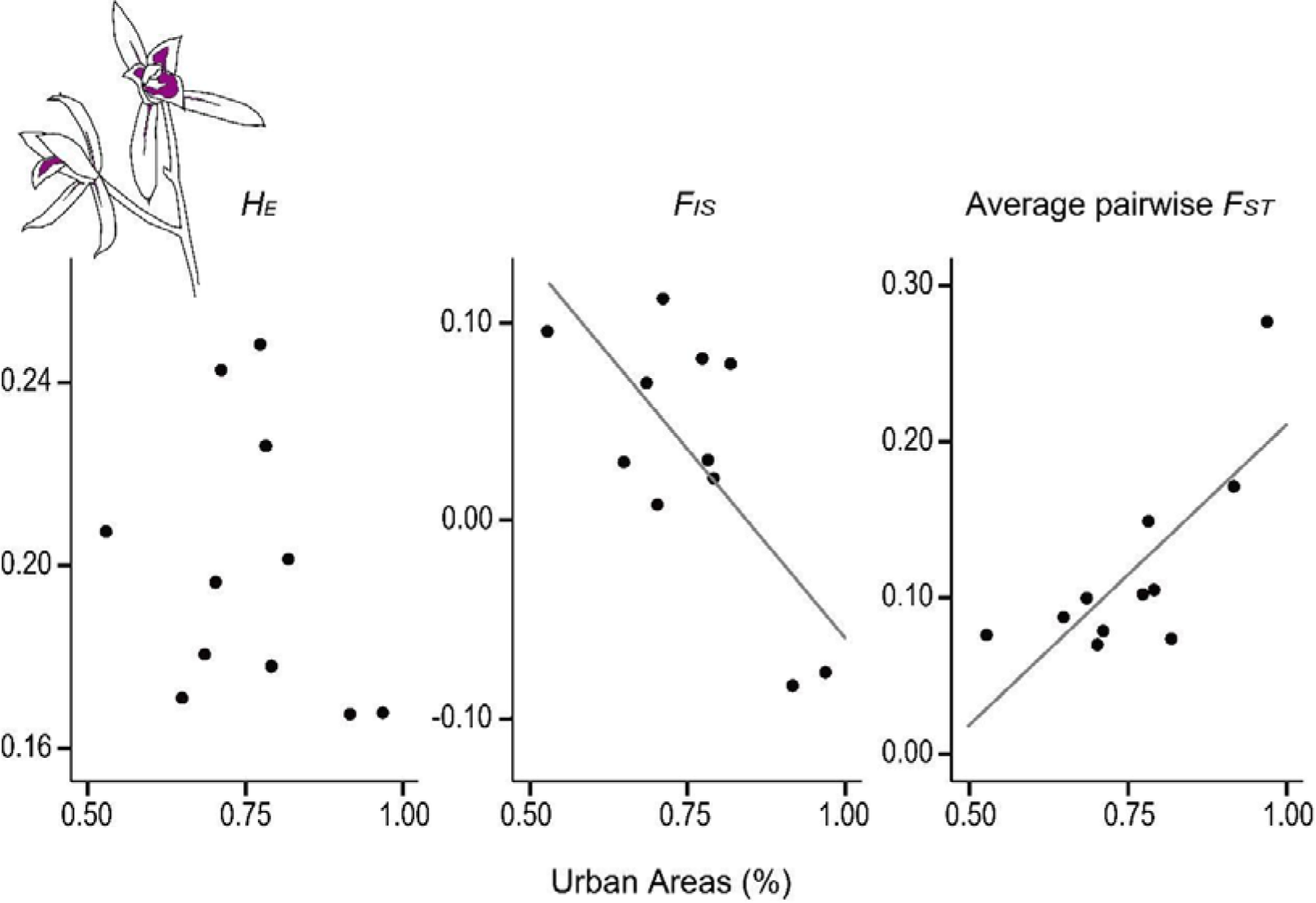
Relationships between the genetic indices and the proportion of urban area within the radius of 5000m in 1976. The significant relationships based on the linear model analysis (in Table 2) were denoted by solid lines.

**Table 2.** Results of the linear model regression analysis, addressing variation of *H*_E_, *F*_IS_, and average pairwise *F*_ST_. Each model included the number of analyzed samples per locus (in Table 1) and the single landscape variable (the proportion of urban/forest area) within a radius of 1000/3000/5000 m as the independent variables. The estimate and 95% confidence interval of the regression coefficient of each landscape variable were shown. The models are ranked according to Akaike’s Information Criterion (AIC), and we calculated the differences in AIC from the best model (ΔAIC).

| Response | Explanatory | | Estimated coefficient | Confidence interval (95%) | | AIC | $\Delta AIC$ | Model Rank |
| --- | --- | --- | --- | --- | --- | --- | --- | --- |
|  | year | area |  | lower | upper |  |  |  |
| $H_E$ | 2020 | Urban 3000 m | -0.175 | -0.400 | 0.051 | -48.294 | 0.000 | 1 |
|  | 1976 | Urban 5000 m | -0.072 | -0.188 | 0.044 | -47.355 | 0.939 | 2 |
|  | 1976 | Urban 3000 m | -0.060 | -0.158 | 0.038 | -47.32 | 0.974 | 3 |
|  | 1976 | Urban 1000 m | -0.031 | -0.110 | 0.048 | -46.301 | 1.993 | 4 |
|  | 2020 | Urban 5000 m | -0.129 | -0.495 | 0.237 | -46.142 | 2.152 | 5 |
| $F_{IS}$ | 1976 | Urban 5000 m | **-0.388** | **-0.595** | **-0.181** | -34.624 | 0.000 | 1 |
|  | 1976 | Urban 3000 m | **-0.308** | **-0.498** | **-0.117** | -32.698 | 1.926 | 2 |
|  | 1976 | Urban 1000 m | **-0.198** | **-0.369** | **-0.027** | -29.228 | 5.396 | 3 |
|  | 2020 | Urban 3000 m | **-0.593** | **-1.143** | **-0.042** | -28.631 | 5.993 | 4 |
|  | 1976 | Forest 5000 m | **0.477** | **0.032** | **0.921** | -28.597 | 6.027 | 5 |
| Average pairwise $F_{ST}$ | 1976 | Urban 5000 m | **0.385** | **0.161** | **0.609** | -32.862 | 0.000 | 1 |
|  | 1976 | Urban 3000 m | **0.292** | **0.078** | **0.505** | -30.207 | 2.655 | 2 |
|  | 1976 | Urban 1000 m | 0.170 | -0.022 | 0.363 | -26.684 | 6.178 | 3 |
|  | 2020 | Urban 5000 m | 0.752 | -0.148 | 1.652 | -26.337 | 6.525 | 4 |
|  | 1976 | Forest 5000 m | -0.405 | -0.900 | 0.090 | -26.221 | 6.641 | 5 |

## Discussion

The present study demonstrated that the low inbreeding coefficients and high genetic differentiation were related to intensive urbanization (mainly the surrounding proportion of urbanized area) at approx. 50 years ago rather than at recent years across the remnant populations of *C. macrorhizon* in the Tokyo MMA.

In this study, the genetic diversity (*H*_E_) of *C. macrorhizon* was the lowest in the highly urbanized area although the linear relationships between genetic diversity and surrounding urban area proportion were not detected. In the plants growing in remnant habitats, the patterns of lower (Hollingsworth and Dickson 1997; Rivkin and Johnson 2022), similar (Culley et al. 2007), and higher (Roberts et al. 2007) level genetic diversity have been reported in urban populations than in rural ones. These patterns, including the result of this study, might indicate that the difference in urbanization impacts on genetic diversity can occur in plant species depending on their life history traits, as in animals (Culley et al. 2007; Suetsugu 2014; Rivkin and Johnson 2022). The mycoheterotrophy in *C. macrorhizon* that mainly specializes the ectomycorrhizal Sebacinales, which closely related to host tree roots (Weiss et al. 2004, 2011; Ogura-Tsujita et al. 2012). This point implies that the presence and population size of host tree species of Sebacinales can regulate the population size of *C. macrorhizon*. The intensive urbanization might lead to the degradation in genetic diversity via both the size reduction and strong isolation of populations of the host tree species of mycorrhizal fungi associated with *C. macrorhizon*, rather than those of forest area.

Although habitat shrinkage and fragmentation by urbanization is expected to result in the high level of inbreeding coefficient (*F*_IS_), which can decrease fitness through inbreeding depression (Reed and Frankham 2003; Lowe et al. 2005), on the contrary highly urbanized populations showed negative inbreeding coefficient, namely heterozygosity excess. Some previous meta-analysis studies have also demonstrated that habitat fragmentations by urbanization and anthropogenic disturbance do not affect inbreeding coefficients significantly and cause heterozygosity excess (Schlaepfer et al. 2018; Miles et al. 2019). Rapid urbanizations in the last century might impose genetic bottlenecks, resulting in heterozygosity excess via genetic drift (Cornuet and Luikart 1996; Stoeckel and Masson 2014; Reynes et al. 2021). Though heterozygosity excess is also explained by specific ecological characteristics such as a lack of selfed progeny, heterosis, negative assortative mating, and asexual reproduction (Stoeckel et al. 2006), these factors have potentials to cause heterozygosity excess in all the studied populations, not in the specific ones. Thus, negative *F*is in highly urbanized sites could be caused by bottleneck impacts. However, we could not exclude the possibility that the extremely small population size in highly urbanized area amplifies the impacts of the ecological characteristics of *C. macrorhizon* on the genetic pattern in a population.

Despite the lack of distinctive population structure based on the individual-level phylogeny across the Tokyo MMA, the genetic differentiation (average pairwise *F*_ST_ between populations) at the population-level increased along with the intensification of urbanization. This pattern supported the prevention of gene flows via habitat isolation due to urbanization (Miles et al. 2019). The high levels of genetic differentiation in the highly urbanized populations based on not only adaptive SNPs but also neutral ones indicated the stochastic changes of genetic composition in each population through genetic drifts associated with population bottlenecks (Toczydlowski and Waller 2019). This result was attributed to the limitation of seed and pollen immigrations by the specificity to the mycorrhizal fungi (Brundrett 2004; Ogura-Tsujita et al. 2012) and the low frequency of insect pollination (Suetsugu 2014). In addition. the habitat condition of *C. macrorhizon* in the remnant forests were not visually changed along the urbanization gradient, but were possibly altered in some ecological aspects (e.g. mycorrhizal or pollinator communities) (Johnson and Munshi-South 2017; Rivkin et al. 2019). Therefore, the stochastic process caused by small population size and modifications of habitat environments in remnant forest floors associated with intense urbanization might lead to both neutral and adaptive divergence of *C. macrorhizon* along the urban-rural landscape.

The present study demonstrated the linear relationships between inbreeding coefficient and genetic differentiation of *C. macrorhizon*, and the past proportion of urbanized area. This pattern implies that the urbanization impacts affected genetic components over at least half a century. This time lag of the alteration in genetic diversity and differentiation, which related to reduced fitness and local extinction, is recognized as the intraspecific-level extinction debt (Helm et al. 2009; Plue et al. 2017; Reisch et al. 2017; Fuller and Doyle 2018; Aavik et al. 2019). The life history of *C. macrorhizon* (e.g. longevity, and dormancy) can produce generation overlaps, which mitigates and prolongs the impacts of habit shrinkage and fragmentation on genetic indices (Ellner and Hairston 1994; Ellner 1996; Tsuzuki et al. 2022). Moreover, the substantial impacts of the relatively expansive landscape within the 5km range on genetic components can be ascribed to the generational overlap, enabling the extended and infrequent dispersal of seeds and/or pollens to genetically rescue a population. The intraspecific-level extinction debt observed in this study partially can contribute to extinction debt of plant species diversity in remnant plant communities of the Tokyo Metropolitan Area. Considering our findings, the substantial extinction debt in global urban areas (Hahs et al. 2009) might be sustained by life history-specific ability to prolong urbanization impacts on genetic diversity in component species.

## Implications for conservation

The present study reported the first evidence for the differences in genetic components due to the delayed impacts of small population size and reduced genetic flow under urbanization. In the 50 years from 1976 to the present, the proportion of urban land-use in the Tokyo MMA have increased by up to 28% (Table S1). Because many species do not become extinct before they are affected by genetic erosion (Spielman et al. 2004), therefore, we should plan to maintain and enhance genetic diversity of remnant species in urban to save these species from local extionctions.

This study did not find substantial genetic differentiation at the individual-level among study sites in the Tokyo MMA. We may enhance the genetic diversity of populations near the center of the MMA by introducing seeds from other populations with the consideration of genetic relatedness. Field seeding techniques were already established for *C. macrorhizon* (Ogura-Tsujita and Yukawa 2008), and the mentioned measures are currently feasible. To ensure the long-term conservation of orchids and other plant species contributing to urban ecosystem services, it is essential not only to assess genetic diversity in urban ecosystems (Gamba and Muchhala 2020; Garner et al. 2020; De Kort et al. 2021) but also to connect its evaluations with life history traits and spatiotemporal impacts of habitat shrinkage and fragmentation due to urbanization (Knapp et al. 2020).

## Acknowledgments

The authors would like to thank Dr. Maiko Kagami for assistance of DNA experiments. This work was financially supported by Grant-in-Aid for Young Scientists (no. 20K20002) and a Grant-in-Aid for Scientific Research B (no. 18H00761 and no. 21H02559) to K.U. from the Ministry of Education, Culture, Sports, Science and Technology of Japan.

## Author’s Contributions

N.I.I. and K.U. performed the sampling, analyses and writing of the manuscript; N.I.I. and S.Y. contributed to the molecular experiments; Y.I. contributed to the field sampling; S.Y. and Y.I. contributed to the writing of manuscript.

## References

Aavik T, Thetloff M, Träger S, et al (2019) Delayed and immediate effects of habitat loss on the genetic diversity of the grassland plant *Trifolium montanum*. Biodivers Conserv 28:3299–3319. 10.1007/s10531-019-01822-8

Brundrett M (2004) Diversity and classification of mycorrhizal associations. Biol Rev Camb Philos Soc 79:473–495. 10.1017/s1464793103006316

Catchen JM, Amores A, Hohenlohe P, et al (2011) Stacks: Building and genotyping loci *de novo* from short-read sequences. G3: Genes Genom Genet 1:171–182. 10.1534/g3.111.000240

Chen S, Zhou Y, Chen Y, Gu J (2018) Fastp: An ultra-fast all-in-one FASTQ preprocessor. Bioinformatics 34:i884–i890. 10.1093/bioinformatics/bty560

Cornuet JM, Luikart G (1996) Description and power analysis of two tests for detecting recent population bottlenecks from allele frequency data. Genetics 144:2001–2014. 10.1093/genetics/144.4.2001

Culley TM, Sbita SJ, Wick A (2007) Population genetic effects of urban habitat fragmentation in the perennial herb *Viola pubescens* (Violaceae) using ISSR markers. Ann Bot 100:91–100. 10.1093/aob/mcm077

De Kort H, Prunier JG, Ducatez S, et al (2021) Life history, climate and biogeography interactively affect worldwide genetic diversity of plant and animal populations. Nat Commun 12:516. 10.1038/s41467-021-20958-2

de Villemereuil P, Gaggiotti OE (2015) A new *F*_ST_-based method to uncover local adaptation using environmental variables. Methods Ecol Evol 6:1248–1258. 10.1111/2041-210X.12418

Dubois J, Cheptou PO (2017) Effects of fragmentation on plant adaptation to urban environments. Philos Trans R Soc 372:20160038. 10.1098/rstb.2016.0038

Duncan RP, Clemants SE, Corlett RT, et al (2011) Plant traits and extinction in urban areas: A meta-analysis of 11 cities. Glob Ecol Biogeogr 20:509–519. 10.1111/j.1466-8238.2010.00633.x

Ellner S (1996) Environmental fluctuations and the maintenance of genetic diversity in age or stage-structured populations. Bull Math Biol 58:103–127. 10.1007/BF02458284

Ellner S, Hairston NG (1994) Role of overlapping generations in maintaining genetic variation in a fluctuating environment. Am Nat 143:403–417. 10.1086/285610

Eriksson O, Kainulainen K (2011) The evolutionary ecology of dust seeds. Perspect Plant Ecol Evol Syst 13:73–87. 10.1016/j.ppees.2011.02.002

Figueiredo L, Krauss J, Steffan-Dewenter I, Sarmento Cabral J (2019) Understanding extinction debts: spatio–temporal scales, mechanisms and a roadmap for future research. Ecography 42:1973–1990. 10.1111/ecog.04740

Fukano Y, Yamori W, Misu H, et al (2023) From green to red: Urban heat stress drives leaf color evolution. Sci Adv 9:eabq3542. 10.1126/sciadv.abq3542

Fuller MR, Doyle MW (2018) Gene flow simulations demonstrate resistance of long-lived species to genetic erosion from habitat fragmentation. Conserv Genet 19:1439–1448. 10.1007/s10592-018-1112-5

Gamba D, Muchhala N (2020) Global patterns of population genetic differentiation in seed plants. Mol Ecol 29:3413–3428. 10.1111/mec.15575

Garner BA, Hoban S, Luikart G (2020) IUCN Red List and the value of integrating genetics. Conserv Genet 21:795–801. 10.1007/s10592-020-01301-6

Goudet J (2005) HIERFSTAT, a package for R to compute and test hierarchical *F*-statistics. Mol Ecol Notes 5:184–186. 10.1111/j.1471-8286.2004.00828.x

Grimm NB, Faeth SH, Golubiewski NE, et al (2008) Global change and the ecology of cities. Science 319:756–760. 10.1126/science.1150195

Hahs AK, McDonnell MJ, McCarthy MA, et al (2009) A global synthesis of plant extinction rates in urban areas. Ecol Lett 12:1165–1173. 10.1111/j.1461-0248.2009.01372.x

Helm A, Oja T, Saar L, et al (2009) Human influence lowers plant genetic diversity in communities with extinction debt. J Ecol 97:1329–1336. 10.1111/j.1365-2745.2009.01572.x

Hollingsworth PM, Dickson JH (1997) Genetic variation in rural and urban populations of *Epipactis helleborine* (L.) Crantz. (Orchidaceae) in Britain. Bot J Linn Soc 123:321–331. 10.1111/j.1095-8339.1997.tb01422.x

Honnay O, Jacquemyn H (2007) Susceptibility of common and rare plant species to the genetic consequences of habitat fragmentation. Conserv Biol 21:823–831. 10.1111/j.1523-1739.2006.00646.x

Huson DH, Bryant D (2006) Application of phylogenetic networks in evolutionary studies. Mol Biol Evol 23:254–267. 10.1093/molbev/msj030

Ives CD, Lentini PE, Threlfall CG, et al (2016) Cities are hotspots for threatened species. Glob Ecol Biogeogr 25:117–126. 10.1111/geb.12404

Japanese Ministry of the Environment (2020) Japanese ministry of the environment red list

Johnson MTJ, Munshi-South J (2017) Evolution of life in urban environments. Science 358:eaam8327. 10.1126/science.aam8327

Johnson MTJ, Prashad CM, Lavoignat M, Saini HS (2018) Contrasting the effects of natural selection, genetic drift and gene flow on urban evolution in white clover (*Trifolium repens*). Proc Royal Soc B 285:20181019. 10.1098/rspb.2018.1019

Kasada M, Matsuba M, Miyashita T (2017) Human interest meets biodiversity hotspots: A new systematic approach for urban ecosystem conservation. PLoS One 12:e0172670. 10.1371/journal.pone.0172670

Khapugin AA, Kuzmin IV, Silaeva TB (2020) Anthropogenic drivers leading to regional extinction of threatened plants: insights from regional Red Data Books of Russia. Biodivers Conserv 29:2765–2777. 10.1007/s10531-020-02000-x

Knapp S, Aronson MFJ, Carpenter E, et al (2020) A research agenda for urban biodiversity in the global extinction crisis. Bioscience 71:268–279. 10.1093/biosci/biaa141

Losdat S, Chang SM, Reid JM (2014) Inbreeding depression in male gametic performance. J Evol Biol 27:992–1011. 10.1111/jeb.12403

Lowe AJ, Boshier D, Ward M, et al (2005) Genetic resource impacts of habitat loss and degradation; reconciling empirical evidence and predicted theory for neotropical trees. Heredity 95:255–273. 10.1038/sj.hdy.6800725

Miles LS, Dyer RJ, Verrelli BC (2018) Urban hubs of connectivity: Contrasting patterns of gene flow within and among cities in the western black widow spider. Proc Royal Soc B 285:20181224. 10.1098/rspb.2018.1224

Miles LS, Rivkin LR, Johnson MTJ, et al (2019) Gene flow and genetic drift in urban environments. Mol Ecol 28:4138–4151. 10.1111/mec.15221

Motomura H, Selosse MA, Martos F, et al (2010) Mycoheterotrophy evolved from mixotrophic ancestors: Evidence in *Cymbidium* (Orchidaceae). Ann Bot 106:573–581. 10.1093/aob/mcq156

O’Grady JJ, Brook BW, Reed DH, et al (2006) Realistic levels of inbreeding depression strongly affect extinction risk in wild populations. Biol Conserv 133:42–51. 10.1016/j.biocon.2006.05.016

Ogura-Tsujita Y, Yokoyama J, Miyoshi K, Yukawa T (2012) Shifts in mycorrhizal fungi during the evolution of autotrophy to mycoheterotrophy in *Cymbidium* (Orchidaceae). Am J Bot 99:1158–1176. 10.3732/ajb.1100464

Ogura-Tsujita Y, Yukawa T (2008) *In situ* seed sowing techniques for the recovery of endangered orchids. Japanese Journal of Conservation Ecology 13:121–127. 10.18960/hozen.13.1_121

Pal R, Meena NK, Pant RP, Dayamma M (2020) *Cymbidium*: Botany, production, and uses. In: Merillon JM, Kodja H (eds) Orchids phytochemistry, biology and horticulture. Springer, Cham, p 4

Peterson BK, Weber JN, Kay EH, et al (2012) Double digest RADseq: An inexpensive method for *de novo* SNP discovery and genotyping in model and non-model species. PLoS One 7:e37135. 10.1371/journal.pone.0037135

Plue J, Vandepitte K, Honnay O, Cousins SAO (2017) Does the seed bank contribute to the build-up of a genetic extinction debt in the grassland perennial *Campanula rotundifolia*? Ann Bot 120:373–385. 10.1093/aob/mcx057

Reed DH, Frankham R (2003) Correlation between fitness and genetic diversity. Conserv Biol 17:230–237. 10.1046/j.1523-1739.2003.01236.x

Reisch C, Schmidkonz S, Meier K, et al (2017) Genetic diversity of calcareous grassland plant species depends on historical landscape configuration. BMC Ecol 17:1–13. 10.1186/s12898-017-0129-9

Reynes L, Thibaut T, Mauger S, et al (2021) Genomic signatures of clonality in the deep water kelp *Laminaria rodriguezii*. Mol Ecol 30:1806–1822. 10.1111/mec.15860

Rivkin LR, Johnson MTJ (2022) The impact of urbanization on outcrossing rate and population genetic variation in the native wildflower, Impatiens capensis. J Urban Ecol 8:juac009. 10.1093/jue/juac009

Rivkin LR, Santangelo JS, Alberti M, et al (2019) A roadmap for urban evolutionary ecology. Evol Appl 12:384–398. 10.1111/eva.12734

Roberts DG, Ayre DJ, Whelan RJ (2007) Urban plants as genetic reservoirs or threats to the integrity of bushland plant populations. Conserv Biol 21:842–852. 10.1111/j.1523-1739.2007.00691.x

Schlaepfer DR, Braschler B, Rusterholz H-P, Baur B (2018) Genetic effects of anthropogenic habitat fragmentation on remnant animal and plant populations: a meta-analysis. Ecosphere 9:e02488. 10.1002/ecs2.2488

Schmidt C, Domaratzki M, Kinnunen RP, et al (2020) Continent-wide effects of urbanization on bird and mammal genetic diversity. Proc Royal Soc B 287:20192497. 10.1098/rspb.2019.2497

Schwarz N, Moretti M, Bugalho MN, et al (2017) Understanding biodiversity-ecosystem service relationships in urban areas: A comprehensive literature review. Ecosyst Serv 27:161–171. 10.1016/j.ecoser.2017.08.014

Seto KC, Güneralp B, Hutyra LR (2012) Global forecasts of urban expansion to 2030 and direct impacts on biodiversity and carbon pools. Proc Natl Acad Sci USA 109:16083– 16088. 10.1073/pnas.1211658109

Shefferson RP (2009) The evolutionary ecology of vegetative dormancy in mature herbaceous perennial plants. J Ecol 97:1000–1009. 10.1111/j.1365-2745.2009.01525.x

Shefferson RP, Jacquemyn H, Kull T, Hutchings MJ (2020) The demography of terrestrial orchids: Life history, population dynamics and conservation. Bot J Linn Soc 192:315–332. 10.1093/botlinnean/boz084

Spielman D, Brook BW, Frankham R (2004) Most species are not driven to extinction before genetic factors impact them. Proc Natl Acad Sci USA 101:15261–15264. 10.1073/pnas.0403809101

Stamatakis A (2014) RAxML version 8: A tool for phylogenetic analysis and post-analysis of large phylogenies. Bioinformatics 30:1312–1313. 10.1093/bioinformatics/btu033

Stoeckel S, Grange J, Fernández-Manjarres JF, et al (2006) Heterozygote excess in a self-incompatible and partially clonal forest tree species — *Prunus avium* L. Mol Ecol 15:2109–2118. 10.1111/j.1365-294X.2006.02926.x

Stoeckel S, Masson J-P (2014) The exact distributions of *F*_IS_ under partial asexuality in small finite populations with mutation. PLoS One 9:e85228. 10.1371/journal.pone.0085228

Suetsugu K (2014) Autonomous self-pollination and insect visitors in partially and fully mycoheterotrophic species of *Cymbidium* (Orchidaceae). J Plant Res 128:115–125. 10.1007/s10265-014-0669-4

Suetsugu K, Ohta T, Tayasu I (2018) Partial mycoheterotrophy in the leafless orchid *Cymbidium macrorhizon*. Am J Bot 105:1595–1600. 10.1002/ajb2.1142

Theodorou P, Albig K, Radzevičiūtė R, et al (2017) The structure of flower visitor networks in relation to pollination across an agricultural to urban gradient. Funct Ecol 31:838–847. 10.1111/1365-2435.12803

Theodorou P, Radzevičiūtė R, Lentendu G, et al (2020) Urban areas as hotspots for bees and pollination but not a panacea for all insects. Nat Commun 11:1–13. 10.1038/s41467-020-14496-6

Toczydlowski RH, Waller DM (2019) Drift happens: Molecular genetic diversity and differentiation among populations of jewelweed (*Impatiens capensis* Meerb.) reflect fragmentation of floodplain forests. Mol Ecol 28:2459–2475. 10.1111/mec.15072

Tsuzuki Y, Sato MP, Matsuo A, et al (2022) Genetic consequences of habitat fragmentation in a perennial plant *Trillium camschatcense* are subjected to its slow-paced life history. Popul Ecol 64:5–18. 10.1002/1438-390x.12093

Uchida K, Karakida K, Iwachido Y, et al (2023) The designation of a historical site to maintain plant diversity in the Tokyo metropolitan region. Urban For Urban Greening 84:127919. 10.1016/j.ufug.2023.127919

Vogt-Schilb H, Munoz F, Richard F, Schatz B (2015) Recent declines and range changes of orchids in Western Europe (France, Belgium and Luxembourg). Biol Conserv 190:133–141. 10.1016/j.biocon.2015.05.002

Wagner CE, Keller I, Wittwer S, et al (2013) Genome-wide RAD sequence data provide unprecedented resolution of species boundaries and relationships in the Lake Victoria cichlid adaptive radiation. Mol Ecol 22:787–798. 10.1111/mec.12023

Weiss M, Selosse M-A, Rexer K-H, et al (2004) *Sebacinales*: a hitherto overlooked cosm of heterobasidiomycetes with a broad mycorrhizal potential. Mycol Res 108:1003–1010. 10.1017/s0953756204000772

Weiss M, Sýkorová Z, Garnica S, et al (2011) Sebacinales everywhere: previously overlooked ubiquitous fungal endophytes. PLoS One 6:e16793. 10.1371/journal.pone.0016793

Willi Y, Van Buskirk J, Hoffmann AA (2006) Limits to the adaptive potential of small populations. Annu Rev Ecol Evol Syst 37:433–458. 10.1146/annurev.ecolsys.37.091305.110145

Yakub M, Tiffin P (2017) Living in the city: urban environments shape the evolution of a native annual plant. Glob Chang Biol 23:2082–2089. 10.1111/gcb.13528

Yukawa T (2015) Orchidaceae. In: Ohashi H, Kadota Y, Murata J (eds) Wild Plants of Japan. Heibonsha, Tokyo, p 193

