## Supplemental information for "Genetic diversity and differentiation of a mycoheterotrophic orchid (*Cymbidium macrorhizon)* under urbanization"

**Supplemental tables and figures**

**TableS1** Locations and landscape variables (the proportion of urban/forest area) within a radius of 1000/3000/5000m of 11 populations of *Cymbidium macrorhizon*.

**TableS2** Moran’s *I* and its significance of genetic indices and landscape variables.

**TableS3** Results of the linear model regression analysis, addressing variation of average pairwise *F*_ST_ based on neutral and adaptive single nucleotide polymorphisms (SNPs). Each model included the number of analyzed samples per locus (in Table 1) and the proportion of urban area within a radius of 5000 m as the independent variables. The estimate and 95% confidence interval of the regression coefficient for the proportion of urban area variable were shown.

**FigureS1** Individual-level phylogenetic relationship (filtered super network) of *Cymbidium macrorhizon* across the Tokyo major metropolitan area (MMA). Filtered super network was estimated based on 1000 maximum likelihood trees with the threshold of 500 trees.

**FigureS2** Population-level phylogenetic relationship (neighbor net) of 11 populations of *Cymbidium macrorhizon* across the Tokyo major metropolitan area (MMA). Neighbor net was estimated based on pairwise *F*_ST_ between the populations.

**FigureS3** Relationships between the average pairwise *F*_ST_ based on neutral/adaptive single nucleotide polymorphisms (SNPs) and the proportion of urban area within the radius of 5000m in 1976. The significant relationships based on the linear model analysis (in Table S2) were denoted by solid lines.

**TableS1**

| Site ID | Lat | Lon | 1976 | | | | | | |  | 2020 | | | | | | |
| --- | --- | --- | --- | --- | --- | --- | --- | --- | --- | --- | --- | --- | --- | --- | --- | --- | --- |
|  |  |  | Urban | | |  | Forest | | |  | Urban | | |  | Forest | | |
|  |  |  | 1000 m | 3000 m | 5000 m |  | 1000 m | 3000 m | 5000 m |  | 1000 m | 3000 m | 5000 m |  | 1000 m | 3000 m | 5000 m |
| 01 | 35°38'22.3" | 139°43'04.1" | 0.923 | 0.973 | 0.917 |  | 0.077 | 0.021 | 0.027 |  | 0.922 | 0.970 | 0.892 |  | 0.070 | 0.018 | 0.036 |
| 02 | 35°40'26.1" | 139°41'38.9" | 0.840 | 0.955 | 0.969 |  | 0.153 | 0.041 | 0.025 |  | 0.640 | 0.923 | 0.951 |  | 0.360 | 0.074 | 0.040 |
| 03 | 35°36'10.8" | 139°35'12.6" | 0.634 | 0.717 | 0.712 |  | 0.249 | 0.117 | 0.114 |  | 0.843 | 0.846 | 0.874 |  | 0.144 | 0.060 | 0.048 |
| 04 | 35°41'56.9" | 139°34'23.6" | 0.852 | 0.852 | 0.819 |  | 0.144 | 0.045 | 0.049 |  | 0.909 | 0.957 | 0.945 |  | 0.075 | 0.016 | 0.027 |
| 05 | 35°36'44.2" | 139°33'44.4" | 0.693 | 0.654 | 0.703 |  | 0.204 | 0.152 | 0.137 |  | 0.705 | 0.824 | 0.865 |  | 0.234 | 0.079 | 0.059 |
| 06 | 35°40'06.6" | 139°33'00.6" | 0.508 | 0.779 | 0.774 |  | 0.148 | 0.065 | 0.049 |  | 0.764 | 0.908 | 0.902 |  | 0.151 | 0.050 | 0.032 |
| 07 | 35°42'31.4" | 139°32'28.3" | 0.780 | 0.825 | 0.792 |  | 0.037 | 0.052 | 0.051 |  | 0.965 | 0.928 | 0.920 |  | 0.007 | 0.039 | 0.034 |
| 08 | 35°23'10.0" | 139°33'13.2" | 0.515 | 0.677 | 0.686 |  | 0.345 | 0.197 | 0.182 |  | 0.652 | 0.867 | 0.846 |  | 0.270 | 0.098 | 0.109 |
| 09 | 35°40'58.7" | 139°31'35.4" | 0.695 | 0.796 | 0.783 |  | 0.214 | 0.073 | 0.050 |  | 0.770 | 0.913 | 0.898 |  | 0.195 | 0.056 | 0.038 |
| 10 | 35°43'19.9" | 139°27'47.6" | 0.510 | 0.632 | 0.650 |  | 0.045 | 0.047 | 0.056 |  | 0.790 | 0.892 | 0.900 |  | 0.009 | 0.018 | 0.024 |
| 11 | 35°33'56.9" | 139°29'08.3" | 0.263 | 0.481 | 0.529 |  | 0.510 | 0.298 | 0.261 |  | 0.558 | 0.764 | 0.812 |  | 0.346 | 0.160 | 0.121 |

**TableS2**

| Variables | | Moran's *I* | *P* value |
| --- | --- | --- | --- |
| *H*_O_ | | 0.026 | 0.222 |
| *H*_E_ | | 0.015 | 0.238 |
| *F*_IS_ | | 0.005 | 0.249 |
| Average pairwise *F*_ST_ | | 0.001 | 0.205 |
| Urban | 1000 m | -0.058 | 0.396 |
|  | 2000 m | **0.273** | 0.009 |
|  | 3000 m | **0.320** | 0.003 |
| Forest | 1000 m | -0.097 | 0.492 |
|  | 2000 m | 0.024 | 0.196 |
|  | 3000 m | **0.176** | 0.030 |

**TableS3**

| Responsible | Explanatory | |  | Estimated coefficient |  | Confidence interval (95%) | |
| --- | --- | --- | --- | --- | --- | --- | --- |
|  | year | area |  |  |  | lower | upper |
| ***Neutral*** SNPs Average pairwise *F*_ST_ | 1976 | Urban 5000 m |  | **0.183** |  | **0.078** | **0.288** |
| ***Adaptive*** SNPs Average pairwise *F*_ST_ | 1976 | Urban 5000 m |  | **0.772** |  | **0.102** | **1.441** |

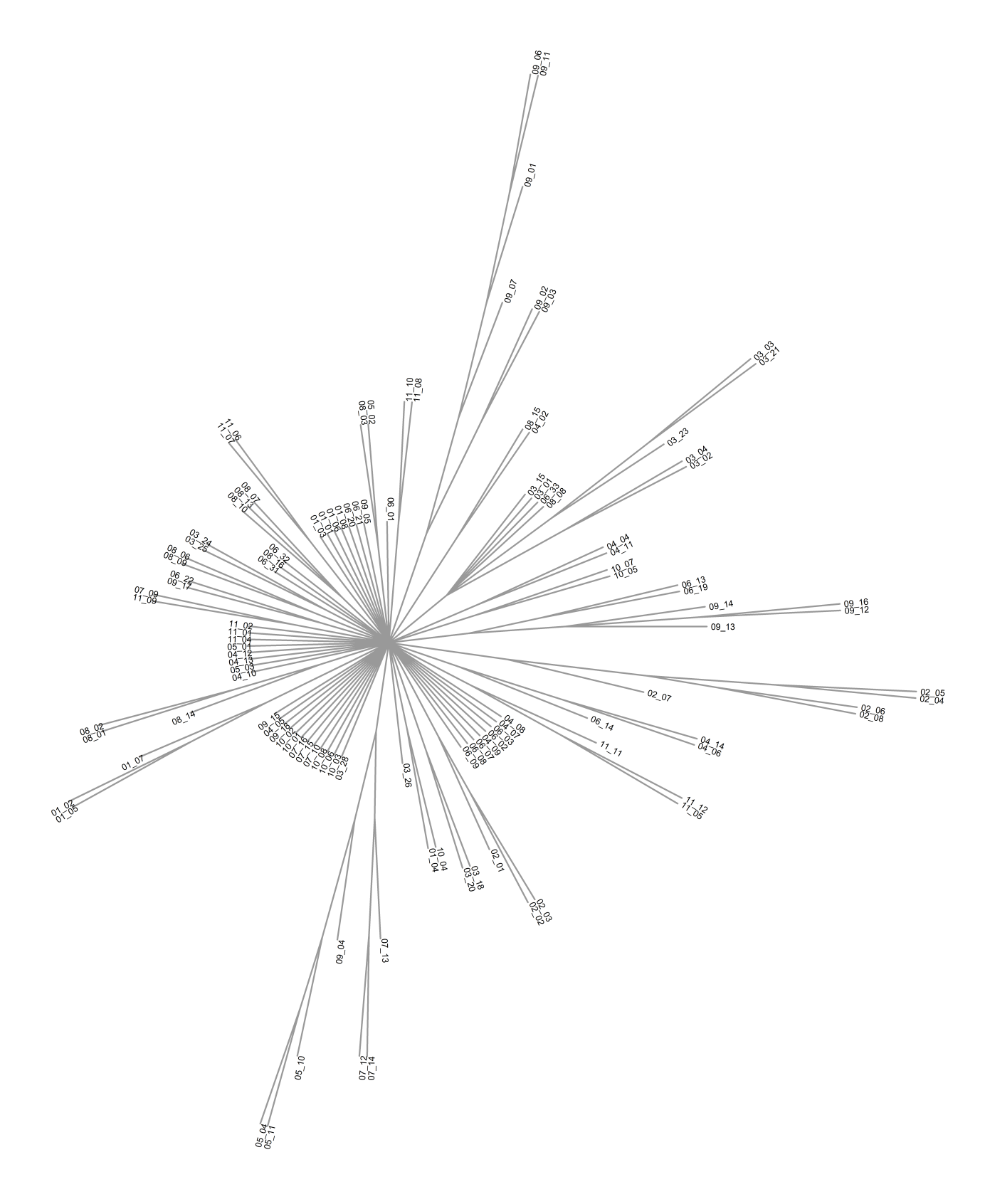
**FigureS1**

**FigureS2**

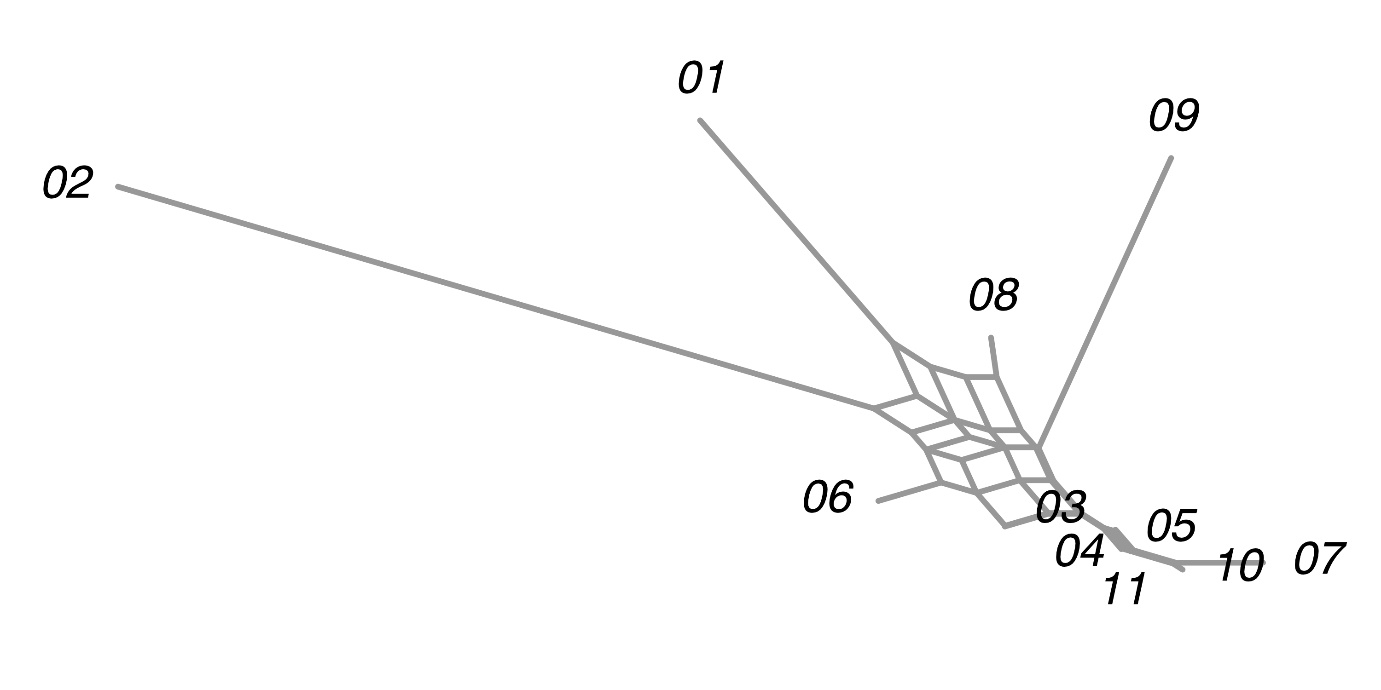

**FigureS3**

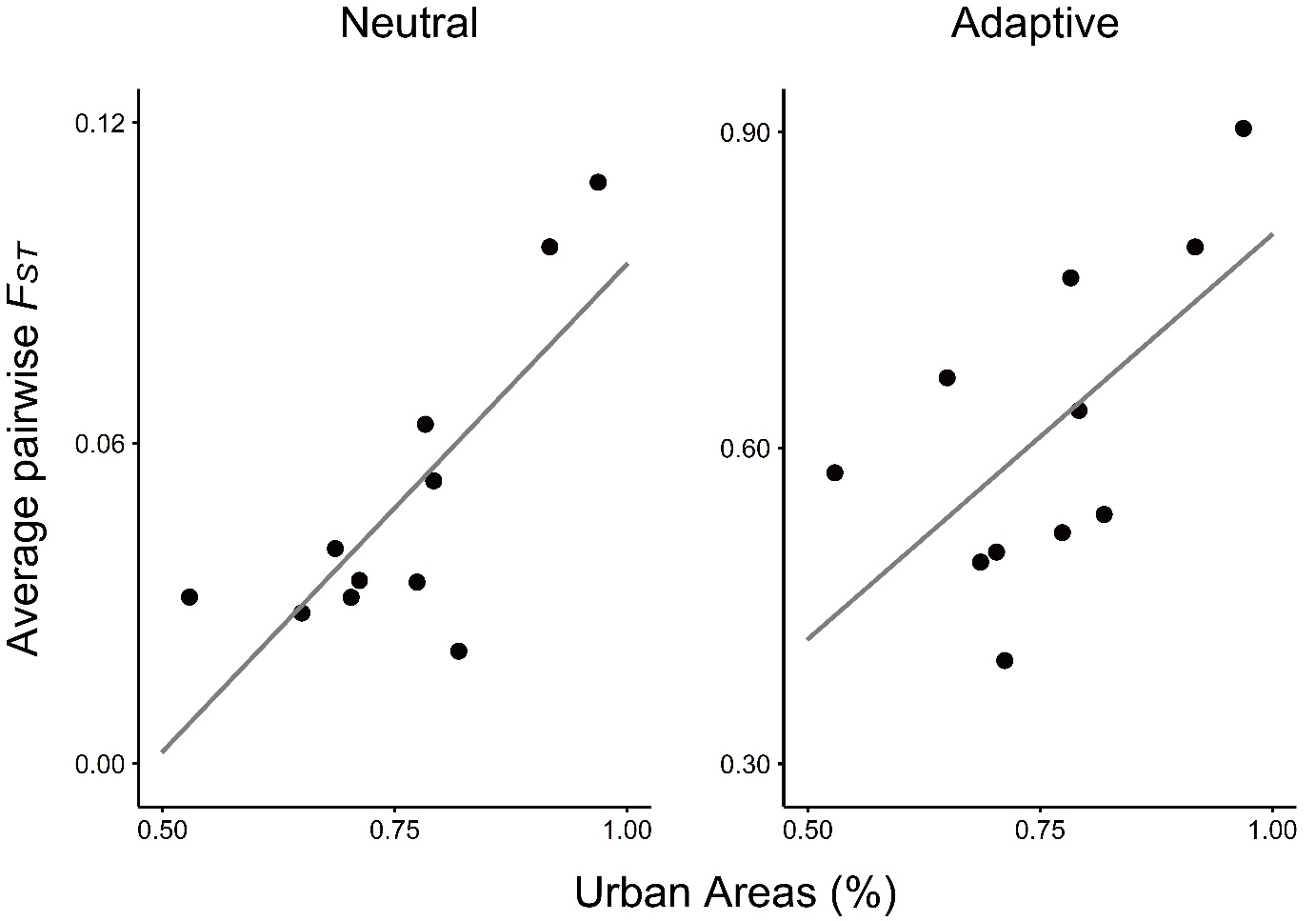
